# Non-Genetic Reprogramming of Monocytes via Microparticle Phagocytosis for Sustained Modulation of Macrophage Phenotype

**DOI:** 10.1101/674598

**Authors:** Kathryn L. Wofford, Bhavani S. Singh, D. Kacy Cullen, Kara L. Spiller

## Abstract

Monocyte-derived macrophages orchestrate tissue regeneration by homing to sites of injury, phagocytosing pathological debris, and stimulating other cell types to repair the tissue. Accordingly, monocytes have been investigated as a translational and potent source for cell therapy, but their utility has been hampered by their rapid acquisition of a pro-inflammatory phenotype in response to the inflammatory injury microenvironment. To overcome this problem, we designed a cell therapy strategy where we collect and exogenously reprogram monocytes by intracellularly loading the cells with biodegradable microparticles containing an anti-inflammatory drug in order to modulate and maintain an anti-inflammatory phenotype over time. To test this concept, poly(lactic-co-glycolic) acid microparticles were loaded with the anti-inflammatory drug dexamethasone (Dex) and administered to primary human monocytes for four hours to facilitate phagocytic uptake. After removal of non-phagocytosed microparticles, microparticle-loaded monocytes differentiated into macrophages and stored the microparticles intracellularly for several weeks *in vitro*, releasing drug into the extracellular environment over time. Cells loaded with intracellular Dex microparticles showed decreased expression and secretion of inflammatory factors even in the presence of pro-inflammatory stimuli up to 7 days after microparticle uptake compared to untreated cells or cells loaded with blank microparticles. This study represents a new strategy for long-term maintenance of anti-inflammatory macrophage phenotype using a translational monocyte-based cell therapy strategy without the use of genetic modification. Because of the ubiquitous nature of monocyte-derived macrophage involvement in pathology and regeneration, this strategy holds potential as a treatment for a vast number of diseases and disorders.

## INTRODUCTION

Tissue regeneration, essential for healing and restoring damaged organs following trauma or disease, is driven by the innate immune system ^1,2^. In particular, monocyte-derived macrophages coordinate multiple aspects of tissue regeneration by homing to sites of injury ^3,4^, phagocytosing pathogenic or necrotic material ^5,6^, secreting factors that drive regeneration ^7,8^, and regulating the behavior of other cells involved in tissue regeneration, including endothelial cells, fibroblasts, and stem cells ^8–13^. Because monocyte-derived macrophages are master regulator cells overseeing tissue regeneration across a number of pathologies, and because monocytes can be readily isolated from peripheral blood, monocytes are an ideal cell source for translational treatments. In fact, the administration of monocytes or macrophages has shown promise in many studies across a range of diseases and tissue injury situations^14^.

First explored in the 1970s as a cancer treatment strategy, monocytes and macrophages have been administered in a number of completed and ongoing clinical trials as a cell therapy for ovarian cancer [NCT02948426], pressure ulcers ^15^, spinal cord injury ^16^, ischemic stroke ^17^, and renal transplant rejections ^18^. In these clinical trials, administration of monocytes or macrophages did not result in severe adverse events, suggesting a high level of safety in administering monocyte/macrophage-based therapies ^14,17^. However, the efficacy of monocyte/macrophage-based therapies has been limited, in large part because macrophages are highly plastic cells that quickly change their behavioral phenotype depending upon microenvironmental stimuli^19^. It is currently not possible to harness the regenerative potential of macrophages because they only transiently induce the desired behavioral pattern before taking on a detrimental phenotype at the site of injury^14,20,21^. For example, when macrophages were polarized to a regenerative phenotype prior to administration into a preclinical model of spinal cord injury in mice, the regenerative macrophages lost this phenotype within 3 days ^20^. Likewise, administration of anti-inflammatory macrophages into a murine model of adriamycin nephropathy lost their anti-inflammatory phenotype within 2 days and upregulated markers of inflammation within 7 days ^21^. In contrast, strategies to control macrophage phenotype via gene editing have shown promise but require further development due to generally non-specific – and potentially detrimental – insertion sites of genetic material into the host genome, low transfection rate, and increased immunogenicity of genetically edited cells ^22,23^. Clearly, there is a need to develop a strategy to promote and sustain an anti-inflammatory phenotype in monocyte-derived macrophages in order to harness their beneficial effects without amplifying local inflammation.

Our overall strategy is to leverage the high therapeutic potential and exceptional behavioral plasticity of monocyte-derived macrophages to mitigate inflammation and promote regeneration following injury. In order to reprogram macrophages and overcome design limitations of previous strategies, such as transient polarization and non-specific targeting, we devised a cell therapy strategy that is selectively administered to monocytes and controls macrophage phenotype over time through the use of intracellular biomaterials. In this strategy, monocytes would be isolated from patients, incubated with drug-loaded microparticles to allow phagocytic uptake, and then re-administered back into the patient either systemically or locally, so that the intracellular release of drug can subsequently modulate inflammation and promote regeneration following injury (Fig. 1). The goal of this study was to characterize the ability of this system to control macrophage phenotype *in vitro* for up to 7 days following microparticle uptake.

**Figure 1.**
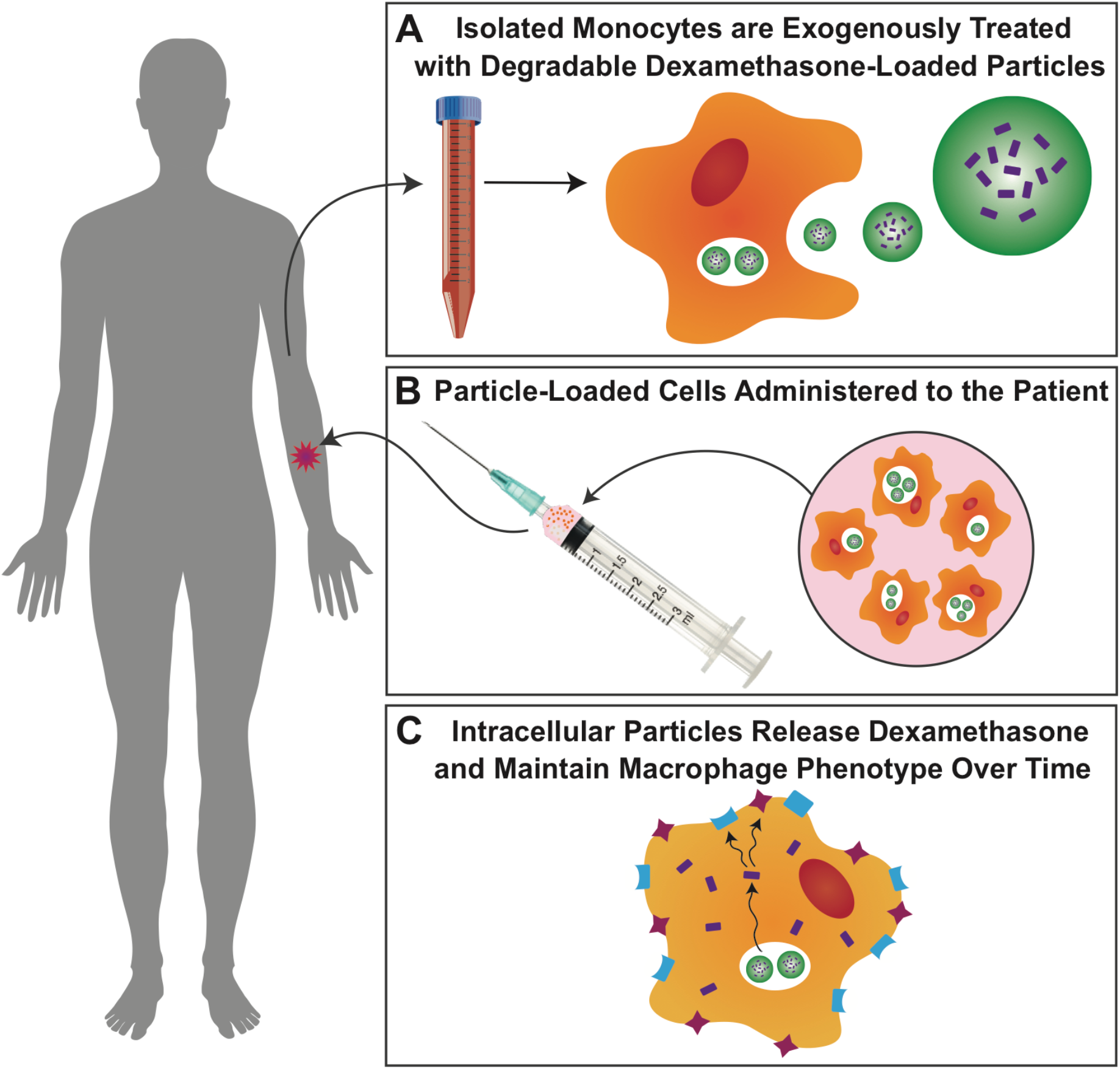
Biomaterial-mediated monocyte cell therapy strategy. (A) Monocytes are isolated from the patient’s blood and incubated with immunomodulatory microparticles, which are rapidly phagocytosed. (B) Microparticle-loaded cells can be administered systemically or locally to the site of injury for a minimally-invasive, autologous treatment. (C) Degradation of the immunomodulatory microparticles over time allows intracellular release of the drug, dexamethasone (Dex), where it can modulate macrophage phenotype.

For proof of concept, we utilized the anti-inflammatory glucocorticoid dexamethasone (Dex) because it down-regulates transcription of inflammatory cytokines ^24–27^ and simultaneously enhances functions associated with regeneration such as homing ^27–29^, phagocytosis ^24,26,27,30,31^, and iron regulation ^27,32,33^ within macrophages. Additionally, Dex receptors are intracellular, making it an ideal choice for an intracellular particle-based strategy. Dex has been widely used for the treatment of inflammatory diseases, but systemic administration affects a broad number of cells ^34–37^, generating deleterious off-target effects that have contributed to a number of failed preclinical and clinical trials investigating Dex as a translational treatment option ^38–40^. Selective delivery of Dex to the monocyte/macrophage population could hold promising therapeutic advantages by avoiding these off-target effects. Previously, we showed that Dex-loaded microparticles could downregulate inflammatory gene expression of primary human macrophages under non-inflammatory conditions *in vitro* ^26,27^. Within this study, we analyzed the ability of the microparticles to release drug intracellularly for several weeks and to maintain macrophage phenotype in the presence of inflammatory stimuli for one week *in vitro*.

## RESULTS

### Microparticle Characteristics and Intracellular Stability

Poly(lactic-co-glycolic) acid (PLGA) microparticles were fabricated with dexamethasone (Dex) or with tetramethylrhodamine (TRITC) as a fluorescent model drug. Microparticles ranged in diameter from 0.98 to 2.05 µm with polydispersity indices between 0.08 and 0.28 (Fig. 2A).

**Figure 2.**
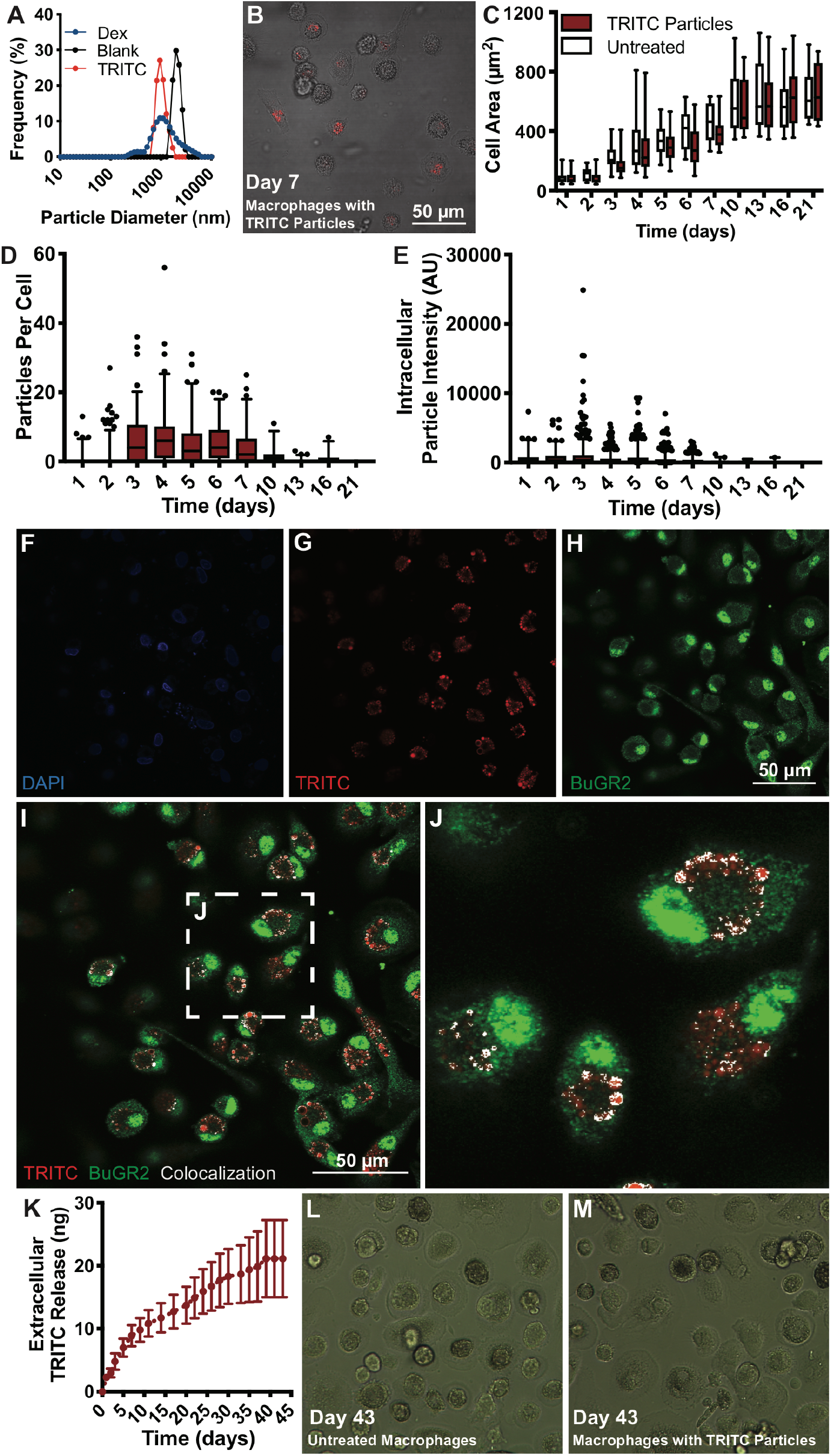
Phagocytosed microparticles release model drugs intracellularly for several weeks. (A) Size distribution of single-emulsion PLGA microparticles fabricated with no drug (Blank), with Dex, or with the fluorescent model drug tetramethylrhodamine (TRITC). (B) Cells loaded with fluorescent microparticles can be imaged over time. (C) Cell area, and thus monocyte-to-macrophage differentiation, is not affected by intracellular microparticle loading. Box and whisker plot represents all data ranging from the minimum to the maximum. (D-E) Intracellular fluorescent microparticles were quantified on a single cell level for number of intracellular microparticles and intracellular microparticle intensity over time per cell (1589 cells analyzed from n=8 experimental replicates). (F-J) Five days after TRITC microparticle administration, cells were stained for nuclei (DAPI, blue) and BuGR2, a glucocorticoid receptor (green) that can be found in the cytoplasm, for imaging along with TRITC (red; n=3). Areas where TRITC signal co-localized with the BuGR2 signal are represented in white. (K) Conditioned media from macrophages loaded with fluorescent microparticles was quantified spectrophotometrically to assess the concentration of extracellular TRITC release from the cells over time (n=22). (L-M) Representative images of untreated macrophages or TRITC microparticle-loaded macrophage morphology and density after 43 days of *in vitro* culture. Data represent mean ± SD for all graphs. Box and whisker plots represent 5-95^th^ percentile of the data with the remaining data plotted as points. Scale bars = 50 µm.

Microparticles were administered to primary human monocytes for 4 hours followed by removal of non-phagocytosed microparticles. Thereafter, monocytes were cultured in macrophage colony stimulating factor (MCSF)-containing media to induce differentiation into macrophages and were imaged at regular intervals (Fig. 2B). Cell area increased over time for both untreated and microparticle-loaded cells, indicating that that monocyte-to-macrophage differentiation was not hindered by intracellular microparticle loading (Fig. 2C). Intracellular fluorescent microparticles were detected for up to 16 days *in vitro* (detection and quantification methods outlined in Sup. Fig. 1). Interestingly, the number of microparticles (Fig. 2D) and their intensity per cell (Fig. 2E) increased in the first three days, which may be due to cell death, phagocytosis of other cells containing microparticles, or microparticle fracture. Thereafter, the intracellular microparticles decreased in number (Fig. 2D) and in fluorescent intensity (Fig. 2E) over time, potentially due to microparticle degradation or reduced fluorophore stability. Staining with BuGR2, a glucocorticoid receptor that can be found in the cytoplasm, indicated that intracellular microparticles were inside the cytoplasm (Fig. 2F-J). Colocalization of TRITC signal and BuGR2 signal suggests that the model drug was intracellularly released and was not confined to the phagolysosomes or endosomes of the cell (Fig. 2I-J). In addition, low levels of TRITC were detected for over 40 days in the extracellular media (Fig. 2K). Because microparticles were rarely observed outside of the cells and because the cell culture media was regularly changed, these results suggest that microparticles remained in the cells at levels that were not detectable via microscopy and released drug into the cell and then extracellularly. Cellular morphology after 43 days of *in vitro* culture was not affected by treatment with TRITC microparticles (Fig. 2L-M).

### Intracellular Dex Microparticles Release Drug Over Time

PLGA microparticles were fabricated with increasing Dex loading, ranging from 0% to 56% w/w Dex (Sup. Fig. 2A). Release of Dex from the 33% and the 56% w/w Dex microparticle groups showed a burst release within the first 24 hours followed by a low but steady release of drug for up to 10 days in PBS (Fig. 3A, Sup. Fig. 2A). Release of Dex from the 2% and 9% w/w Dex microparticle groups exhibited no burst release but exhibited a slow, steady release of drug for more than one week in PBS (Fig. 3A, Sup. Fig. 2A). To characterize extracellular Dex release from macrophages containing intracellular microparticles, cells were treated with microparticles prepared with 33% and 56% w/w Dex microparticles. We detected low levels of Dex in the extracellular space of microparticle-loaded cells on days 3, 5, and 7 (Fig. 3B, Sup. Fig. 2B).

**Figure 3.**
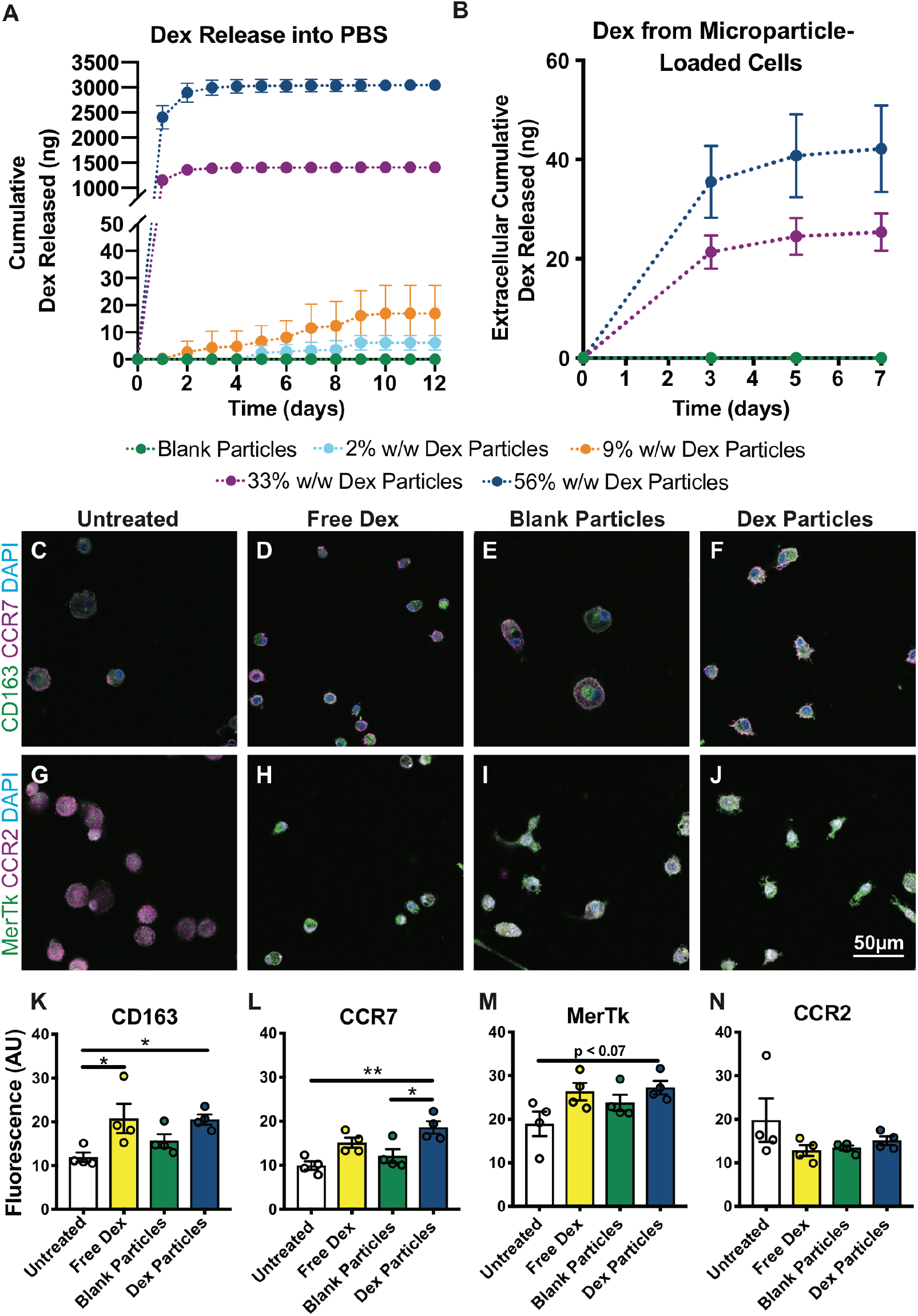
Intracellular Dex microparticles modulate cell behavior one week after microparticle treatment. (A) Release profile of Dex from microparticles in PBS (n=5). (B) Dex content in the cell culture media (n=3 in blank group and n=6 in treatment groups) following Dex microparticle administration to cells. Media was changed on days 3, 5, and 7. (C-J) Representative images of cells stained for CD163 and CCR7 or stained for MerTk and CCR2. (K-N) Image quantification of staining for the surface receptors CD163, CCR7, MerTk, and CCR2 (n=4). Data represent mean ± SEM for all graphs. Statistical analyses were completed by applying one-way ANOVA with Tukey’s post hoc test. *denotes p<0.05 and ** denotes p<0.01. Scale bar = 50 µm.

### Intracellular Dex Microparticles Modulate and Maintain Macrophage Phenotype

We next examined the effects of Dex microparticles on expression of receptors associated with phagocytosis and homing, which are critical behaviors in macrophage-mediated regeneration. At 7 days after microparticle administration, microparticle-loaded cells upregulated expression of CD163, a surface receptor associated with hemoglobin-haptaglobin scavenging during tissue regeneration ^41–43^, compared to untreated controls, reaching similar levels as those observed in cells treated continuously with free Dex (Fig. 3C; Sup. Fig. 3). Likewise, CCR7, a surface receptor associated with chemotaxis through damaged tissues ^44,45^, was upregulated in cells treated with Dex microparticles relative to untreated controls and cells treated with blank microparticles (Fig. 3D; Sup. Fig. 3). MerTK, which is associated with phagocytosis of apoptotic cells and cellular debris ^30,46^, was modestly increased in Dex microparticle-treated cells compared to untreated controls (p<0.07). Contrary to our expectation, we did not observe a significant increase in expression of CCR2, a homing-associated receptor, in cells treated with free Dex or Dex microparticles (Fig. 3E-F) even though previous literature has shown that Dex upregulates this receptor in macrophages ^26,28,30^.

### Intracellular Dex Microparticles Modulate Macrophage Phenotype in the Presence of Pro-Inflammatory Stimuli

Because numerous pathologies are characterized by prolonged inflammation, we next wanted to investigate if the intracellular microparticles could maintain macrophage phenotype even in the presence of inflammatory stimuli. We expected that since Dex was released intracellularly, it could bind to intracellular glucocorticoid receptors and inhibit pro-inflammatory pathways even when cells were cultured in inflammatory environments. To test this, we administered microparticles with increasing Dex concentrations to monocytes and cultured the particle-loaded cells in inflammatory conditions (complete media supplemented with lipopolysaccharide (LPS) and interferon gamma (IFNγ)) for 7 days. We observed that Dex microparticles caused decreases in the secretion of tumor necrosis factor alpha (TNFα) in a dose-dependent manner (Fig. 4A-B). Furthermore, cells treated with microparticles containing high levels of Dex behaved similarly to cells that were continuously treated with free Dex (Fig. 4A-B).

**Figure 4.**
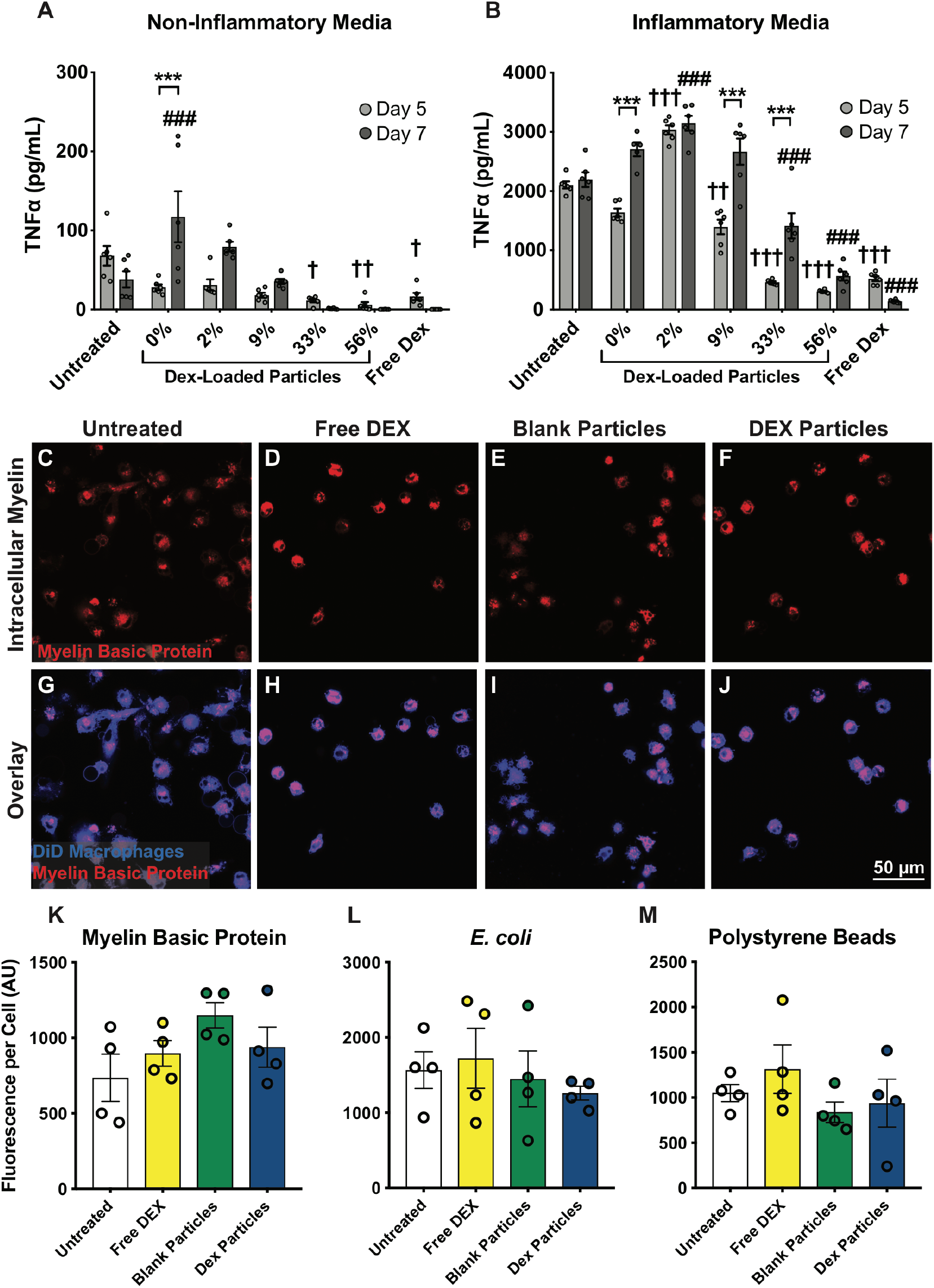
Intracellular Dex microparticles modulate cell behavior even in the presence of inflammatory stimuli. Tumor necrosis factor alpha (TNFα) protein secretion was measured in conditioned media from microparticle-treated cells in (A) non-inflammatory or (B) inflammatory media (n=6). Data represent mean ± SEM. Two-way ANOVAs with Tukey’s post hoc tests were completed using p values corrected for multiple comparisons. *denotes p<0.05, **denotes p<0.01, ***denotes p<0.001, ^**†**^denotes significant differences relative to untreated controls on day 5, and ^#^denotes significant differences relative to untreated controls on day 7. (C-J) Representative images of DiD-labeled monocytes that were untreated, treated with continuous free Dex, treated with blank microparticles, or treated with Dex microparticles for four hours prior to removal of non-phagocytosed microparticles, and then cultured in inflammatory media conditions for 5 days prior to incubation with phagocytosis targets. Phagocytic uptake of (K) myelin basic protein, (L) *E. coli*, or (M) polystyrene beads within six hours was quantified across treatment condition (between 6 and 27 cells analyzed and averaged from each experimental replicate with n=4 experimental replicates). Data represent mean ± SEM for all graphs. Statistical analyses were completed by applying one-way ANOVA with Tukey’s post hoc analysis with corrected p values. No statistical differences were detected. Scale bar = 50 µm.

### Intracellular Dex Microparticles Do Not Preclude Phagocytosis

Previous research suggests that Dex administration to macrophages can enhance phagocytosis ^24,47^, but we were curious if pre-loading monocytes with intracellular microparticles would prevent any subsequent phagocytosis. We completed an *in vitro* functional assay of phagocytosis in the presence of inflammatory stimuli using three different types of targets that an immune cell might encounter: recombinant myelin basic protein, a difficult protein to digest that contributes to pathology following nervous system injury ^48,49^; *E. coli*, a common bacterium that macrophages might eliminate from sites of infection ^50^; and carboxylated polystyrene beads, a common biomaterial that could represent phagocytosis of foreign debris. Microparticle-loaded cells were cultured in inflammatory conditions with the phagocytosis targets for six hours on day 5. Pre-loading cells with microparticles did not prevent subsequent phagocytosis of any of the three targets compared to untreated controls (Fig. 4C-M).

### Intracellular Dex Microparticles Modulate Inflammation, Phagocytosis, and Homing Genes

We next examined the effects of Dex microparticles on transcription of genes related to inflammation, phagocytosis, homing, and iron metabolism because these genes are modulated by Dex and/or are essential for tissue regeneration ^24,26,47^. Gene expression data were first assessed with a heatmap where both the rows (representing individual genes) and the columns (representing experimental replicates) were organized with hierarchical clustering (Fig. 5A). The six experimental replicates within each treatment condition clustered together for all four treatment groups. Cells treated with Dex microparticles were most closely associated with cells that were treated with continuous free Dex (Fig. 5A). Additionally, cells treated with blank microparticles were most closely associated with the negative control cells that were untreated.

**Figure 5.**
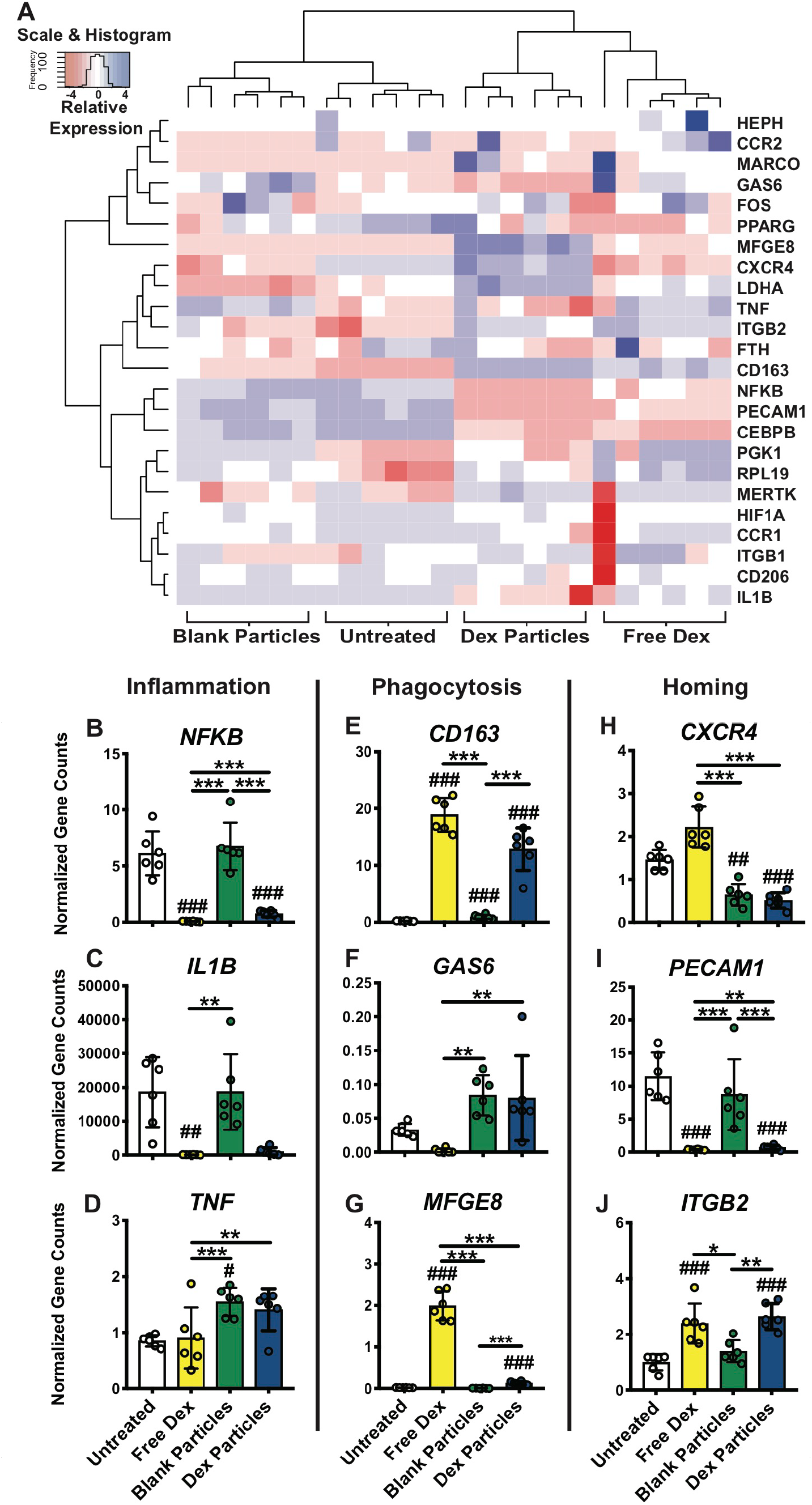
Intracellular microparticles modulate macrophage gene expression. (A) Gene expression, processed with a reciprocal transform, was plotted in a heatmap with scaling by rows. Dendrogram organization of the heatmap columns (samples across treatment conditions) was employed to organize samples according to similar gene expression profiles (n=6). A subset of genes related to (B-D) inflammation, (E-G) phagocytosis, and (H-J) homing were plotted. All genes analyzed can be found in the supplementary figures 4 and 5. One-way ANOVA statistical analyses were completed on transformed data from each of the data sets. To facilitate interpretation, non-transformed data were plotted in column graphs. Tukey’s post hoc analyses with corrections for multiple comparisons were completed as appropriate. *denotes p<0.05, **denotes p<0.01, ***denotes p<0.001, and ^#^denotes significant differences relative to untreated controls.

Dex microparticles caused macrophages to downregulate genes associated with inflammation, including *NFKB*, *IL1B*, (Fig. 5B-D; Sup. Fig. 4A-D). However, unlike the protein secretion data (Fig. 4A-B), *TNF* gene transcription was increased in both the blank microparticle group and the Dex microparticle group, suggesting that post-transcriptional modification could have influenced TNFα protein secretion levels. In agreement with studies describing the effects of Dex on macrophage gene expression^31,42^, genes related to phagocytosis such as *CD163* and *MFGE8* were significantly upregulated in the Dex microparticle group relative to untreated controls (Fig. 5E, G; Sup. Fig 4E-K). *GAS6*, a gene associated with phagocytosis, was upregulated in both the blank microparticle group and the Dex microparticle group relative to the free Dex control group (Fig. 5F). Considering the published effects of Dex on monocyte homing behavior ^28,29^, we were surprised to find that most of the homing-related genes were not up-regulated in response to free Dex or Dex microparticles (Fig. 5H-J). Five of the six homing genes investigated were not significantly different or were downregulated in groups treated with Dex microparticles relative to untreated controls (Sup. Fig. 5A-F). Both the blank microparticle group and the Dex microparticle group downregulated the homing gene *CXCR4* (Fig. 5H). In contrast, the free Dex control group and the Dex microparticle group down-regulated *PECAM1* expression but up-regulated *ITGB2* expression relative to untreated controls and blank microparticle controls (Fig. 5I-J). Although previous studies have shown that Dex can modulate some functions associated with iron metabolism ^33^ and other studies have found that different macrophage phenotypes regulate environmental iron in distinct ways ^51^, we observed that administration of free continuous Dex or Dex microparticles did not increase genes related to iron metabolism (Sup. Fig. 5G-J). In fact, expression of *CEPBP*, which encodes the CEBPB transcription factor that has been associated with immune and inflammatory responses, was decreased in cells treated with free Dex or Dex microparticles.

## DISCUSSION

The behavioral plasticity of monocyte-derived macrophages allows these cells to play many crucial roles in tissue regeneration, but also poses a challenge in cell therapy because they fail to maintain desirable phenotypes upon administration. In this study, we have demonstrated proof of concept that loading monocytes with drug-loaded biodegradable microparticles can modulate the phenotype of macrophages derived from those monocytes for up to 7 days *in vitro*. To our knowledge, this study represents the first report of reprogramming of monocytes for sustained maintenance of macrophage phenotype without genetic modification. Taken together with studies that have shown that macrophages can deliver drugs to sites of interest ^52,53^, our findings suggest that microparticle-loaded monocytes hold considerable potential as a cell therapy strategy across a wide range of applications.

In recent years, several landmark studies have shown that the release of immunomodulatory drugs from implanted biomaterials can modulate the behavior of host monocytes and macrophages, with beneficial effects in treating disease and tissue injury ^11,54–57^. For example, the release of the S1P receptor agonist FTY720 recruits regenerative monocytes and macrophages to promote tissue regeneration ^57–59^. However, the release of drug from biomaterials may affect all cells in the vicinity, not just macrophages, which may result in off-target effects. Other studies have targeted monocytes by injecting drug delivery vehicles into the blood stream, because circulating monocytes are highly phagocytic ^60^. However, in this strategy the majority of nanoparticles and microparticles become sequestered in the liver, lung, spleen, and kidneys, resulting in wasted drug and the potential for side effects in these organs ^61^. In addition, the uptake of microparticles by circulating monocytes can inhibit their ability to home to the site of injury, redirecting them to the spleen and causing apoptosis ^61^. By loading monocytes with immunomodulatory biomaterials *ex vivo* and re-administering locally to sites of injury, the disadvantages of off-target effects and reduced homing could be mitigated.

In the present study we showed that some homing genes were downregulated by Dex microparticle treatment while others were upregulated, but these results are likely confounded by the fact that we analyzed the cells after their differentiation into macrophages, which is known to inhibit homing capacity relative to monocytes ^62^. Nonetheless, it is likely that the microparticles would need to be optimized to maintain homing capabilities of monocytes in order for this cell therapy strategy to be utilized as a systemic administration.

Previous literature found that administration of indigestible particles did not prevent macrophage phagocytosis of *E. coli* or polystyrene beads ^63^. Likewise, within this study, we observed that pre-loading cells with Dex microparticles did not preclude any subsequent phagocytosis of targets such as myelin basic protein, *E. coli*, or polystyrene beads. In the context of gene expression, administering blank microparticles or Dex microparticles generated similar results in several genes, such as *CXCR4*, *TNF*, and *GAS6*, perhaps suggesting that the act of phagocytosing or storing a microparticle intracellularly can modulate macrophage gene expression even in the absence of a therapeutic compound.

The results of this study suggest that intracellular Dex microparticles can modulate and maintain anti-inflammatory macrophage phenotype over time. However, this study is not without limitations. Most importantly, our goal was to assess the capacity of the intracellular microparticles to maintain an anti-inflammatory phenotype of macrophages after 7 days in a highly inflammatory environment *in vitro*, but the therapeutic benefits of this approach remain to be demonstrated with *in vivo* models. In particular, the dose of drug, number of microparticles, number of macrophages, and timing of administration will need to be optimized in order to assess a therapeutic effect. Moreover, we chose dexamethasone for proof of concept, but a drug that promotes a more regenerative macrophage phenotype would be a better choice for studying therapeutic efficacy. Finally, within this paper we have not characterized the effect of microparticle-loading on the homing capacity of monocytes, although other groups have completed similar characterizations ^52,53,62^, but this will be important if monocytes are to be administered systemically.

In conclusion, we demonstrated proof of concept that Dex-loaded microparticles, administered to monocytes, could act intracellularly to modulate and maintain the phenotype of monocyte-derived macrophages for up to one week *in vitro*. Importantly, this methodology could be utilized as a minimally-invasive, autologous treatment method for a variety of injuries and diseases. Further understanding and advancing tools that modulate immune cell behavior will be instrumental in overcoming detrimental pathology and promoting tissue regeneration across a number of organ systems and disease states.

## ACKNOWLEDGEMENTS

This work was supported by the National Institutes of Health (R01-HL130037 to KLS); the Department of Veteran’s Affairs (RR&D Merit Review #I01-RX001097 to DKC); and the Department of Education GAANN iCARE Fellowship (to KLW).

## METHODS

### Microparticle Fabrication

Single emulsion poly(lactic-co-glycolic) acid (PLGA) microparticles were fabricated by dissolving PLGA (Lakeshore Biomaterials; cat. #5050 DLG4A) at 20 mg/mL in an organic solution containing 9:1 dichloromethane (DCM; Acros Organics, cat. #364230010) to trifluoroethanol (TFE; Sigma-Aldrich, cat. #T63002). Depending on the group, 100 µg/mL of the model drug tetramethylrhodamine (TRITC; Invitrogen; cat. #A1318), 400 µg/mL of the model drug Nile Red (Acros Organics; cat. # 415711000), and/or increasing concentrations of dexamethasone (Dex; Alfa Aesar, cat. #A17590) ranging from 0 to 25 mg/mL were added to the organic phase. Blank microparticles were fabricated in the absence of any therapeutic or model drugs. Thereafter, the organic phase was suspended in 2% poly(vinyl alcohol) (PVA; Sigma Aldrich, cat # 363170) and sonicated for 60 seconds on ice. The mixed solution was added to an even larger volume of PVA and allowed to stir for 6 hours to facilitate solvent evaporation and microparticle hardening. Following solvent evaporation, microparticles were centrifuged at 4300rpm for 10 minutes, washed in DI H_2_O, sonicated again for 30 seconds on ice to separate any microparticle aggregates. Size and polydispersity index of microparticles were quantified via dynamic light scattering on a Malvern Zetasizer. Microparticles were frozen, lyophilized, sterilized with UV light, resuspended at 1 mg/mL in sterile 1x phosphate buffered saline (PBS), and stored at −80°C until use.

### Primary Human Monocyte Cell Culture

Human primary monocytes were purchased from the University of Pennsylvania’s Human Immunology Core. Monocytes were initially frozen at 10*10^6^ cells/mL in freezing media containing 50% RPMI-1640 media (Glibco, cat. #11875-093), 40% heat inactivated human serum (HIHS; Sigma, cat. #H3667), and 10% dimethyl sulfoxide (DMSO, Fisher Scientific, cat. #BP231-100). For standard cultures with human primary monocytes, cells were thawed, freezing media was removed, and cells were suspended at 2*10^6^ cells/mL in media (89% RPMI-1640, 10% HIHS, and 1% penicillin-streptomycin (Pen/Strep; Glibco, cat. #15140-122)) supplemented with the appropriate treatment. Cells were treated with: 20 µg of microparticles for every million cells; 39.25 µg/mL of Dex (free Dex control); or with nothing (untreated control). Cells were allowed to incubate for 4 hours at 37°C while shaking gently. Thereafter, cells were centrifuged down at 400xg for 7 minutes, the supernatant containing non-phagocytosed microparticles was removed, and cells were plated at 1*10^6^ cells/mL in ultra-low attachment wells for experiments related to protein secretion, gene expression, or extracellular drug release or were plated in glass-bottom chamber slides for experiments related to imaging. All conditioned media was supplemented with 20 ng/mL of macrophage colony stimulating factor (MCSF; Peprotech, cat. #300-25) to induce monocyte to macrophage differentiation. Cell media in the “Free Dex” group was further supplemented with 39.25 µg/mL Dex at every media change. Media was completely removed and replaced on day 3, 5 and 7. To generate an inflammatory environment, complete cell media was supplemented with 100 ng/mL of lipopolysaccharide (LPS; Sigma, cat. #L2654) and 100 ng/mL of interferon gamma (IFNγ; Peprotech, cat. #300-02) on days 3, 5, and 7.

### Characterizing Fluorescent Microparticle Localization

In order to characterize intracellular microparticle stability, fluorescent microparticles containing the model drugs Nile Red or TRITC were administered to human primary monocytes. Thereafter, monocytes were cultured on glass chamber slides, incubated at 37°C with media changes every 2 to 3 days. Live cells were imaged with confocal microscopy every 1-3 days. In order to identify microparticle distribution within the cells, a subset of wells was treated with TRITC microparticles on day 0, and then washed, fixed, and stained immunohistochemically on day 5 to investigate the extent of TRITC colocalization with the glucocorticoid receptor BuGR2.

### Immunocytochemistry

Fluorescent immunocytochemistry was employed to determine cytoplasmic and surface marker distribution and intensity. Either on day five or day seven, cell media was removed, and cells were washed twice in 1x PBS. Washed cells were fixed in 10% paraformaldehyde (Electron Microscopy Science, cat. #15710-S) at room temperature for 20 minutes and then washed in 1x PBS three more times. Washed cells were incubated in blocking solution containing 5% normal horse serum (Invitrogen, cat. # 31874) and 0.1% triton-X (Sigma, cat. #X-100) for one hour at room temperature. Blocked cells were then incubated overnight at 4°C with primary antibodies against BuGR2 (Invitrogen, cat. #MA1-510), CD163 (1:50; MyBioSource, cat. # MBS302638), CCR7 (1:100; OriGene, cat. # TA320232), MerTk (1:200; abcam, cat. # ab216564), and/or CCR2 (1:200; abcam, cat. # ab176390). Cells were washed and then incubated with the appropriate secondary antibodies for 4 hours at room temperature against anti-rabbit (1:100; Invitrogen, cat. #A-11034), anti-mouse (1:100; abcam, cat. #ab150119), or anti-rat (1:100; Invitrogen, cat. #A-21247). Sections were also counterstained with 4′,6-diamidino-2-phenylindole (DAPI; 1:1000; Invitrogen, cat. #D1306). Immunocytochemical images were acquired with a confocal microscope (Olympus FV1000 Laser Scanning Confocal) using fluorescence or differential interference contrast (DIC) with the 20x or 60x objectives.

### Image Analysis of Confocal Micrographs

In order to systematically process confocal micrographs, we generated a custom MATLAB code to quantify the number and intensity of intracellular microparticles in a set of images (Sup. Fig. 1). Briefly, DIC and fluorescent channels were separated so that cell and microparticle channels could be processed independently. The DIC channel, containing information about the cells, was converted to a grayscale image and regions of high contrast were identified. These regions of interest were then closed by dilating and eroding the edges. Small noise was eliminated, holes within each object were filled, clusters of cells were removed, and objects on the border of the image were removed (Sup. Fig. 1). The remaining segmented regions of interest were considered a representative population of cells within the image. This segmentation was validated by overlaying the segmentation overtop of the original raw image. The number of pixels in each object in the cell segmentation mask was utilized to determine cell area.

Separately, the fluorescent channel was processed but analysis varied slightly depending on the experiment. For the intracellular microparticle longevity study, the fluorescent channel represented intracellular microparticles, so the channel was quantified for number of particles, particle fluorescent intensity, and total cellular fluorescent intensity (a measure of both the number of particles and particle intensity). For the immunocytochemistry study, the fluorescent channels represented cell protein distribution, so the fluorescent channels were quantified for intensity within the cell segmentation.

Particle quantification in the particle longevity study was completed by converting the particle fluorescent channel to binary and applying a watershed transform to separate clusters of particles (Sup. Fig. 1). Only particles within the regions of the cell segmentation were considered. This eliminated the occasional extracellular particle and only considered intracellular particles within single cells not adjacent to the image border. The particle segmentation was validated by overlaying the particle segmentation on top of the original raw image. Thereafter, the number of intracellular microparticles and the intracellular microparticle intensity was quantified within each individual cell segmentation. This generated a rich dataset where we could assess how individual cell size, microparticle number within each cell, and microparticle fluorescent intensity within each cell changed over time. Immunocytochemistry fluorescence was completed by quantifying the total image fluorescence that was within the regions of the cell segmentation. This metric normalized fluorescence to the total cell area within each image. This process was completed separately for each fluorophore of interest.

### Characterizing Dex Release Kinetics

Dex release from microparticles was completed to assess release kinetics over time. 40µg of microparticles containing Dex were suspended in 1x PBS inside tubes containing a 200 nm membrane (Pall Corporation, cat. #ODM02C33). The microparticles were incubated at 37°C while shaking gently at 100 rpm. At regular intervals, the microparticle-containing solution was separated from microparticles by centrifugation at 200xg for 5 minutes. The flow-thru solution was collected and quantified spectrophotometrically for absorbance at 241 nm, and the microparticles were resuspended in fresh 1x PBS (Sup. Fig. 2). A Dex standard curve was made at 241 nm in warmed 1x PBS to correlate absorbance with concentration.

### Enzyme-Linked Immunosorbent Assays

At regular intervals, conditioned media collected from cells was assessed with enzyme-linked immunosorbent assays (ELISA) in order to quantify the concentration of extracellular tumor necrosis factor alpha (TNFα, Invitrogen, cat. #88-7346-88) or Dex (Bioo Scientific, cat. #1112). All ELISAs were completed according to the manufacturer’s instructions.

### Phagocytosis Assay

Human primary monocytes were fluorescently labeled with Vybrant DiO cell-labeling solutions (Molecular Probes, cat. #V-22886) according to the manufacturer’s instructions. Thereafter, fluorescent monocytes were incubated with nothing, with free Dex, with blank microparticles, or with Dex microparticles for 4 hours at 37°C while shaking. Following incubation, cells were separated from non-phagocytosed microparticles and cultured in regular media in a glass chamber slide. Media was supplemented with free Dex in appropriate conditions. On day 3, media was removed and replaced with inflammatory media and with free Dex in appropriate conditions. On day 5, either human recombinant myelin basic protein (Alfa Aesar, cat. #AAJ66051LB0) fluorescently labeled with Vybrant DiD (Molecular Probes, cat. # 22887), fluorescent *Escherichia coli* (*E. coli*) BioParticles (ThermoFisher, cat. #E2863), or fluorescent carboxylate-modified 1 µm polystyrene beads (Invitrogen, cat. #F8816) were added to cell cultures suspended in inflammatory media. The final target concentrations were 50 µg/mL for myelin basic protein, 1*10^7^ particles/mL of *E. coli*, and 1*10^7^ particles/mL for the polystyrene beads. Cells were allowed to incubate with the fluorescent targets at 37°C for 6 hours. Following incubation, media was removed, cells were washed three times, and were fixed in 10% paraformaldehyde at room temperature for 20 minutes. Cells were washed two more times and then imaged on a confocal microscope (Olympus FV1000 Laser Scanning Confocal). In order to measure phagocytic uptake, images were computationally segmented, as described above, and the total fluorescence signal per cell was quantified.

### RNA Extraction

Primary human monocytes were thawed and treated with nothing, free Dex, blank microparticles, or Dex microparticles and allowed to mix gently at 37°C for four hours. Following incubation, cells were separated from non-phagocytosed microparticles, resuspended in complete media supplemented with MCSF, and plated in ultra-low attachment plates. On days 3 and 5, media was gently removed and replaced with complete media with MCSF supplemented with inflammatory stimuli LPS and IFNγ. On day 7, media was removed, and lysis buffer was added to each well. Cell lysates were stored at −80°C until RNA purification could be completed. RNA from human cells was cleaned and collected utilizing an RNAqueous-Micro Kit (Ambion, cat. #AM1931) according to the manufacturer’s instructions. RNA concentration was quantified using a NanoDrop 1000 Spectrophotometer (V 3.8.1).

### Gene Expression Analysis

Gene expression analysis was completed by using NanoString’s high-throughput PlexSet technology to assess absolute gene counts from purified RNA. A custom code set was generated and corresponding oligonucleotides probes, which bind to specific genes of interested, were fabricated through Integrated DNA Technologies. Thereafter, experimental samples, oligonucleotide probes, and NanoString reagent were added together according to the manufacturer’s instructions. Absolute gene count raw data was quantified with the nCounter MAX prep station. Raw gene counts were first normalized by subtracting out the lane-spectific negative controls and then dividing by the lane-specific positive controls. Negative values were set to zero and data was transformed according to a reciprocal transform *y’ = 1/(y+1)*. The heat map presents transformed data and all statistics were completed on transformed data because transformation generated a normal distribution. However, to ease interpretation for the readers, column graphs were plotted of data prior to transformation because a reciprocal transform function would present data in a counter-intuitive manner.

### Statistical Analyses

All data were statistically assessed with the appropriate parametric test. If we observed that data did not adhere to parametric data requirements, data was transformed to generate an approximately normal distribution and subsequently characterized with the appropriate parametric one-way or two-way ANOVA. P values from Tukey’s Honest Significant Difference post hoc analyses were always adjusted to account for multiple comparisons. Data generated from NanoString quantification was assessed with a one-way ANOVA for each gene on transformed data. However, to simplify data interpretation, the pre-transformed data were plotted within each gene’s graph.

