## Supplemental Figures for "Non-Genetic Reprogramming of Monocytes via Microparticle Phagocytosis for Sustained Modulation of Macrophage Phenotype"

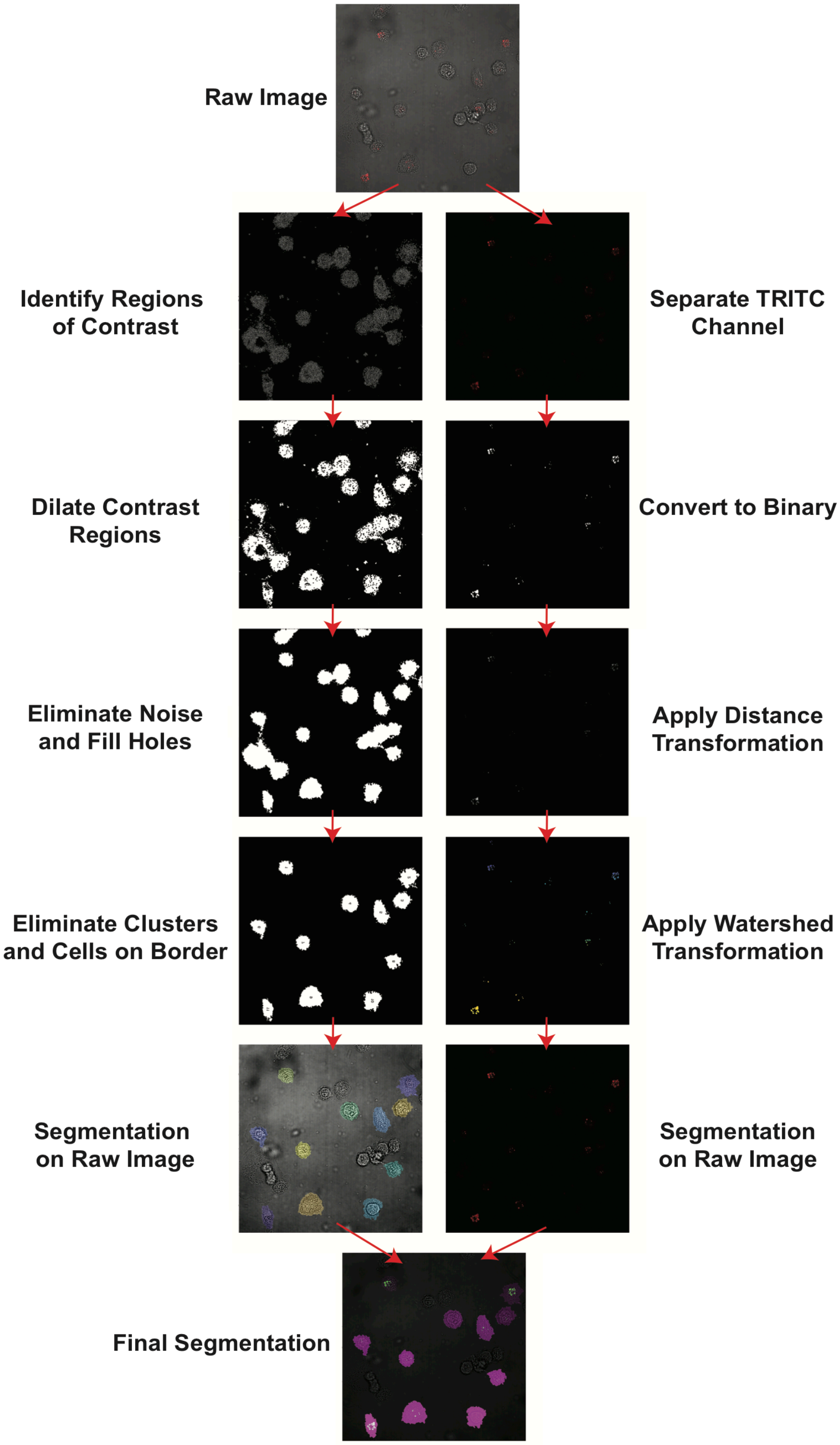

Raw Image
Identify Regions of Contrast
Separate TRITC Channel
Dilate Contrast Regions
Convert to Binary
Eliminate Noise and Fill Holes
Apply Distance Transformation
Eliminate Clusters and Cells on Border
Apply Watershed Transformation
Segmentation on Raw Image
Segmentation on Raw Image
Final Segmentation

**Supplementary Figure 1. Flow chart of feature-based segmentation for image analysis.** Confocal images from live cell cultures treated with TRITC microparticles were quantified over time with a custom-built MATLAB code. First, the TRITC channel and the cell channel were separated. Then, regions of contrast were identified and dilated in the cell channel. Next, noise within the binary mask was eliminated, holes within larger objects were filled, and cells in clusters or on the image border were eliminated to complete the cell channel segmentation. Separately, the TRITC channel was converted to binary and a Watershed transform was applied in order to separate adjacent microparticles. The final segmentation of both channels was overlaid on the original, raw image to ensure accurate cell and microparticle segmentation. Thereafter, data from each cell (including cell area, number of microparticles in each cell, and intracellular microparticle intensity within each cell) was extracted.

**A**

| | Dex in organic phase (μg/mL) | PLGA in organic phase (μg/mL) | Dex in the organic phase (% w/w) | Dex released from 40μg of particles (ng) | Percent weight from cumulative release (%) |
|---|---|---|---|---|---|
| Blank Particles | 0 | 20 | 0.0 | 0.0 | 0.000 |
| 2% Dex-loaded particles | 0.4 | 20 | 2.0 | 6.1 | 0.006 |
| 9% Dex-loaded particles | 2 | 20 | 9.1 | 16.9 | 0.017 |
| 33% Dex-loaded particles | 10 | 20 | 33.3 | 1406.9 | 1.407 |
| 56% Dex-loaded particles | 25 | 20 | 55.6 | 3043.0 | 3.043 |

**B**

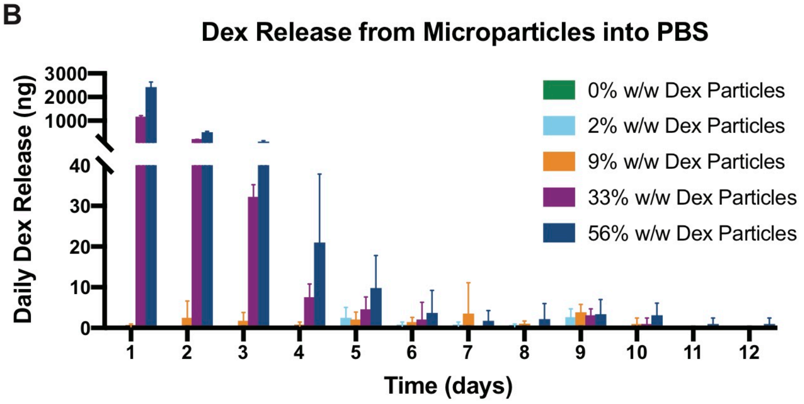


**C**

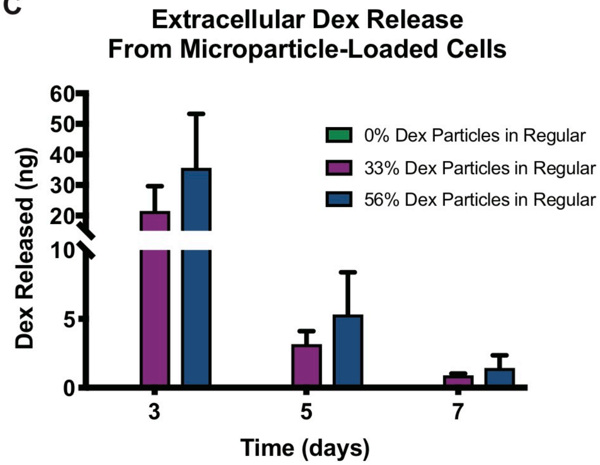


**Supplementary Figure 2. Daily release of Dex from microparticles.** (A) Table of fabrication parameters for Dex-loaded particles. (B) Quantity of Dex released into PBS from Dex-loaded microparticles at each time point. (C) Quantity of Dex released into the extracellular space from macrophages loaded with Dex microparticles at each time point. Data represent the mean ± SD.

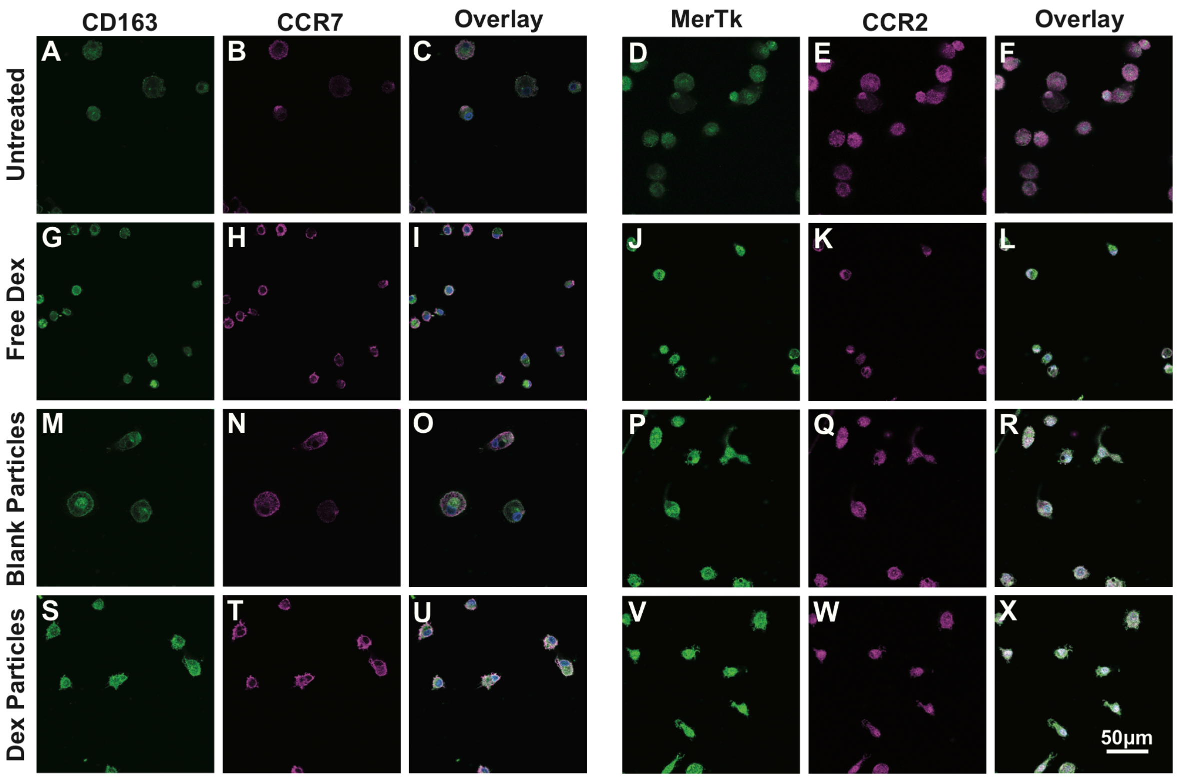


**Supplementary Figure 3. Intracellular Dex microparticles increase surface receptor expression seven days after microparticle treatment.** Monocyte were untreated (A-F), treated with continuous free Dex (G-L), loaded with blank microparticles (M-R), or loaded with Dex microparticles (S-X). Representative images of staining for the surface receptors CD163, CCR7, MerTk, and CCR2 (n=4).

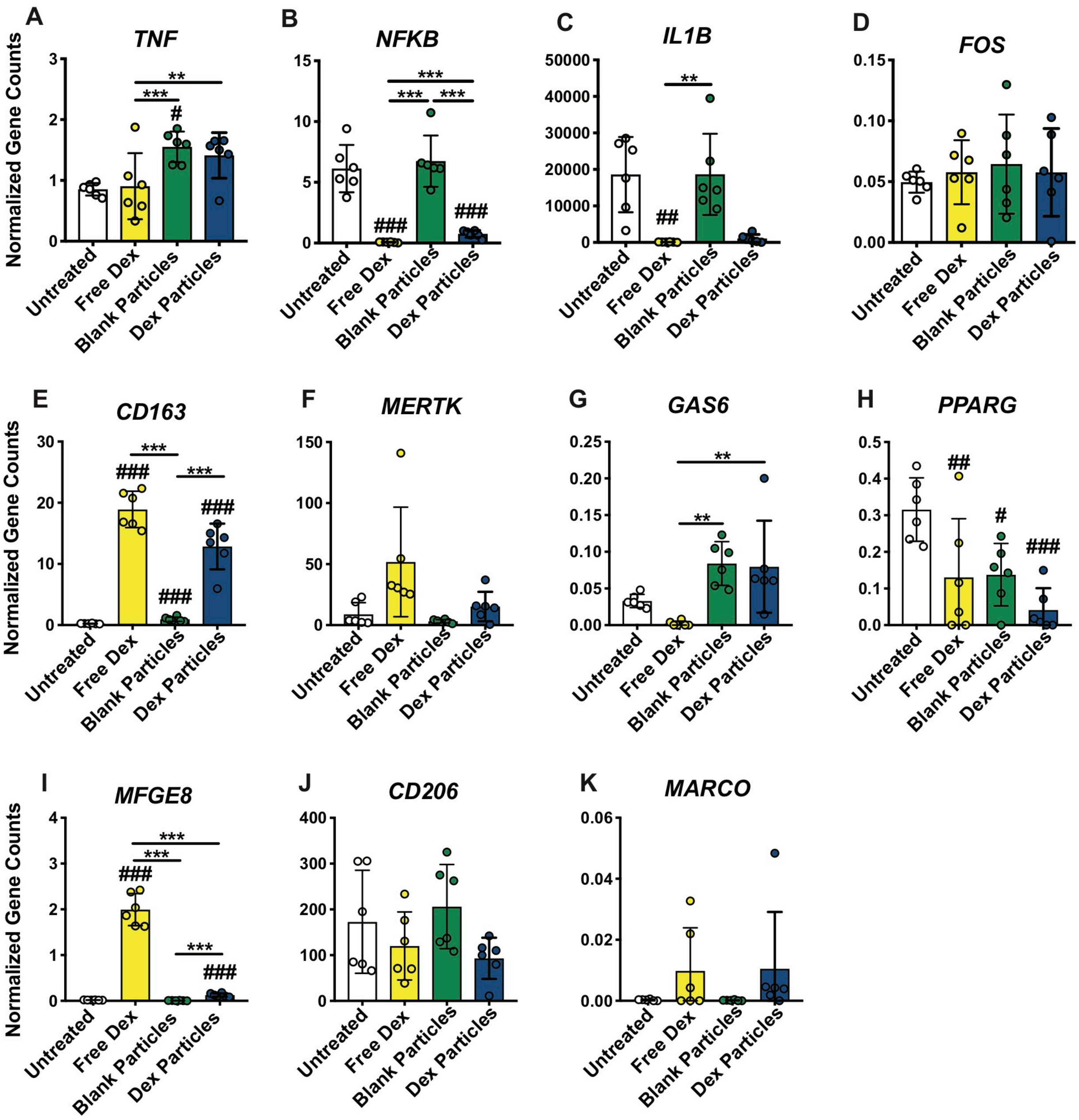


**Supplementary Figure 4. Intracellular microparticles modulate macrophage gene expression related to inflammation and phagocytosis**. Human primary monocytes that were untreated, treated with continuous free Dex, treated with blank microparticles, or treated with Dex microparticles were cultured in inflammatory medial for seven days and then analyzed for gene expression (n=6). Genes related to (A-D) inflammation or (E-K) phagocytosis were plotted. One-way ANOVA statistical analyses were completed on transformed data from each of the data sets. Tukey's post hoc analyses with corrections for multiple comparisons were completed on appropriate data. *denotes $p<0.05$, **denotes $p<0.01$, ***denotes $p<0.001$, and $^{\#}$denotes significant differences relative to untreated controls.

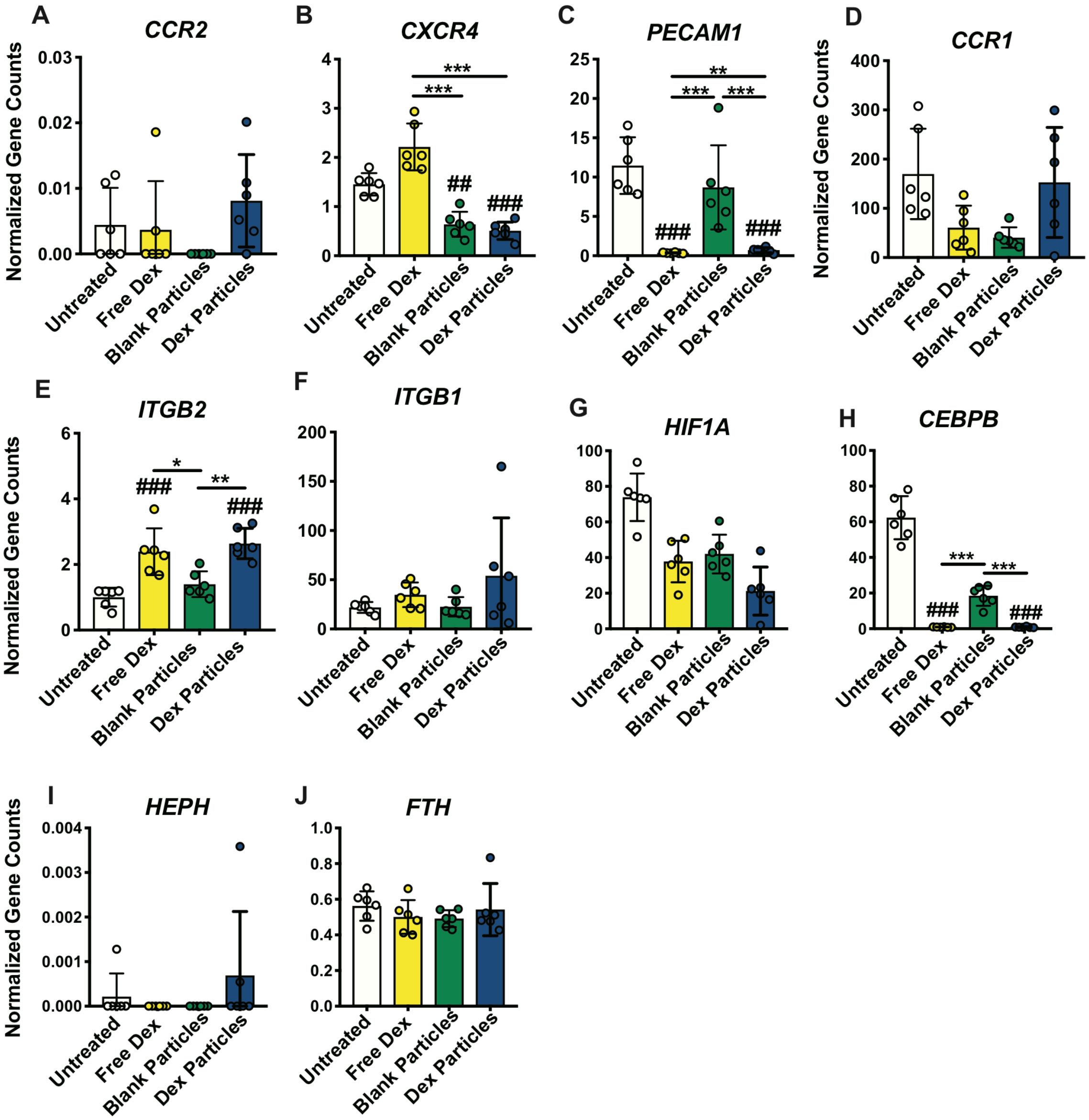


**Supplementary Figure 5. Intracellular microparticles modulate macrophage gene expression related to homing and iron metabolism**. Human primary monocytes that were untreated, treated with continuous free Dex, treated with blank microparticles, or treated with Dex microparticles were cultured in inflammatory media for seven days and then analyzed for gene expression (n=6). Genes related to (A-F) homing or (G-J) iron metabolism were plotted. One-way ANOVA statistical analyses were completed on transformed data from each of the data sets. Tukey's post hoc analyses with corrections for multiple comparisons were completed on appropriate data. *denotes $p<0.05$, **denotes $p<0.01$, ***denotes $p<0.001$, and #denotes significant differences relative to untreated controls.
